# Cutting Edge: An mRNA Platform to Create Isolated, Monospecific Th1 Responses

**DOI:** 10.1101/2025.07.23.666239

**Authors:** Kathleen S. Krauss, Stephen Carro, Michael J. Hogan, Laurence C. Eisenlohr

## Abstract

Helper T cells (T_CD4_) are lynchpins of adaptive immune responses. Since each T_CD4_ expresses a single T cell receptor, recognizing an epitope within major histocompatibility complex class II (MHCII), it is often desirable to study a population that responds to the same epitope. We devised a novel method of selective immunization to produce a robust, monospecific T_CD4_ in mice using a modular mRNA vector. The vector encodes the target epitope attached via flexible linker to MHCII. Immunization with this mRNA selectively elicits T_CD4_ responses across a range of epitopes and MHCII alleles. These T_CD4_ show robust, polyfunctional Th1 cytokine release when evaluated *in vitro*. Additionally, we tested the activity of these T_CD4_ cells during *Salmonella enterica* infection, a model of Th1-dependent immune response, and demonstrated their efficacy *in vivo* in curtailing infection.

## Introduction

Helper T cells (T_CD4_) are a critical component of the adaptive immune response. T_CD4_ are required for 1) release of cytokines to coordinate immune responses (1–4); 2) development of diverse, high affinity antibody responses (5–7); 3) durable B cell and cytotoxic T cell memory responses (8–12). These effects are mediated by a diverse repertoire of T_CD4_, each of which responds to a specific epitope presented by MHC class II (MHCII), often by distinct pathways and antigen-presenting cell types (13–16). This diversity within such a crucial cell subset has provoked substantial interest in studying the impact of CD4 T cell responses to individual epitopes.

Within the T_CD4_ repertoire there are a variety of subtypes; our inquiries focus on Th1 cells, one of the first to be identified and one of the best characterized T_CD4_ phenotypes. Th1s play a crucial role in immune responses to intracellular bacteria and viruses, by both providing help to other lymphocytes and exhibiting direct lysis of infected cells. To carry out these activities, Th1s produce a signature array of cytokines: IFNγ, TNFα, and IL-2 (17, 18). The critical role of Th1s in immune responses have been demonstrated in archetypal models of infection such as *Salmonella enterica*, where depletion of the cells themselves or their signature cytokines have severe consequences (19–25); conversely, immunizing mice with a specific CD4 epitope can offer protection against lethal infection (26).

To study the effect of epitope specificity *in vivo*, one must produce a population of monospecific CD4 T cell response in animals. There are two general approaches to achieve this: adoptive transfer and immunization. Adoptive transfer requires the isolation of T_CD4_ from a donor mouse, typically a transgenic mouse line which expresses only one T cell receptor (TCR); these T cells are transferred into recipient mice of the same genetic background (27, 28). Advantages of adoptive transfer include precise control over number of cells administered and straightforward tracking of cells via congenic surface markers. However, a significant limitation of this approach is the need for existing TCR transgenic mouse lines, which are expensive and time-consuming to produce.

The second general strategy to produce epitope-specific CD4 responses is peptide immunization. Soluble peptide can be used to directly immunize against a single epitope; however, this strategy triggers a broad immune response beyond T_CD4_ and frequently employs inflammatory adjuvants such as complete Freund’s adjuvant, whose use has been discouraged due to animal welfare concerns (29–33). One can also isolate antigen presenting cells (APCs), saturate their surface MHC with soluble peptide, and infuse these cells into mice. Such peptide-pulsed APCs have been well-validated for provoking T_CD8_ or mixed T cell responses (34–42). However, when it comes to eliciting T_CD4_ cells specifically, this method has only been cited in a model employing MHCII knockout recipient mice (43). It is not clear why peptide-pulsed DCs are so unreliable for generating T_CD4_ responses.

Here we present a system to induce robust, epitope-specific T_CD4_ in mice using an mRNA construct encoding an epitope linked to the N-terminus of the MHC-II beta chain (pMHCIIβ). Our platform is modular, allowing linkage of any epitope by oligo cloning into the mRNA-encoding plasmid. Synthesis and delivery of the mRNA is possible with commercially available materials, including an “off-the-shelf” lipid nanoparticle (LNP) mRNA encapsulation reagent, which makes it accessible to labs without prior experience in mRNA/LNP production. Like clinical mRNA vaccines, our platform can be easily modified to update the encoded sequence, and it provokes a strong T cell response without a requirement for adjuvants beyond the LNP, which is well tolerated. In addition, the *Salmonella enterica* infection model demonstrates the robust functionality of these cells *in vivo*.

## Materials and Methods

### mRNA preparation

mRNA preparation was as previously described (46–48). Briefly, endotoxin-free preparations of the plasmid (available upon request) encoding mRNA for pMHCIIβ were linearized via restriction digest, purified via phenol/chloroform/isoamyl alcohol extraction, then used to synthesize mRNA with the Invitrogen MEGAscript™ T7 Transcription Kit, substituting N1-Methylpseudouridine-Triphosphate for the uridine triphosphate supplied in the kit and adding the TriLink CleanCap® reagent at 1mM. mRNA was purified with cellulose to remove ds RNA products.(46) Concentration and purity were verified via Nanodrop 2000.

### In vitro mRNA transfection and T hybridoma activation assay

mRNA preparations encoding pMHCIIβ were complexed using the Mirus TransIT mRNA transfection kit and delivered to HEK293T (ATCC) or immortalized fibroblasts from either C57BL/6 or BALB/c mice stably transfected to express CIITA.(49) Cells had been passaged the day prior and plated at 5e4 cells/well in triplicate in a 96-well, flat bottom, black plate. After transfection, T hybridomas(50) were added at 10^5^ cells/well. Cells were cultured for 16-20h, then antigen presentation was quantified as described previously.(16)

### Mice and immunizations

Wildtype female C57Bl/6J mice (strain 000664) were purchased from the Jackson Laboratory and maintained under SPF conditions at the Children’s Hospital of Philadelphia, under an IACUC-approved protocol. mRNA-LNPs were prepared using the JetRNA+ kit by Polyplus according to the manufacturer protocols and administered intraperitoneally on d0 and d6 to mice 6-8 weeks of age.

### Characterization of T cell responses *ex vivo*

10-12 days after the first immunization, mice were euthanized by CO_2_ inhalation and spleens were collected. Splenocytes were suspended after RBC depletion via ACK lysis, then cultured with antigen at a final concentration of 10μg/mL and anti-CD28 antibody (Tonbo biosciences) at a final concentration of 8ng/mL. For secreted proteome assay, culture media was isolated by centrifugation after 24h, then analyzed using Proteome Profiler Mouse Cytokine Array Kit from Biotechne. For intracellular cytokine staining, cells were treated with Brefeldin A 30min after antigen addition. After 12h, cells were stained with the following: Live/Dead Fixable Aqua (Invitrogen); Fc block (BD Biosciences); BD Cytofix/Cytoperm Kit; and CD8-PerCP/Cy5.5, CD3-PacBlue, CD4-BV786, IFNg-AF488, and either TNFa-BV650 and IL2-PE (eBioscience) or CCL3-APC and GMCSF-PE (Invitrogen). Flow cytometry data was acquired with a Cytoflex LX (Beckman) and analyzed with FlowJo software (Tree Star).

### ELISA characterization of serum antibody

Serum was collected post-mortem from mice immunized with pMHCIIβ targeting Ova_323_/I-A^b^, then analyzed via ELISA. Briefly, ELISA plates were coated with either Ova_323_ peptide from Genscript or Ova_329-337_/I-A^b^ monomer from the NIH tetramer core, then incubated with serum samples followed by anti-mouse detection antibody (62-6520) and SeraCare TMB Chromogenic Kit.

### Infection of mice with Salmonella enterica strains

Immunized C57Bl/6 mice were infected with 5×10^8^ CFU of *Salmonella enterica* serovar Typhimurium SL7207 (hisG46, DEL407 [aroA544::Tn10 (Tcs)]) expressing truncated MisL protein with 4 repeats of Ova_323_ at the N terminus from an AmpR plasmid, a kind gift from Dieter Schifferli, University of Pennsylvania.(51) Overnight liquid cultures were diluted to 50^9^/mL in LB (estimated by OD_600_ and confirmed by serial dilution plating); 100μL was administered to mice via gavage. 1dpi and 4dpi fresh stool was collected, weighed, and suspended in PBS. Samples were cultured on LB-amp plates to determine CFU.

### Statistical analysis

Statistical analysis was performed using GraphPad Prism 10. *p* values <0.05 were considered statistically significant.

## Results and Discussion

### A customizable mRNA transcript allows immunization targeted to a specific epitope/MCHII allele pair

We have designed a modifiable mRNA transcript to allow selective immunization against a single T_CD4_ epitope. This transcript encodes the target epitope at the N terminus attached to the MHCIIβ allele of choice via a flexible linker of (GGGGS)3, similar in design to that designed by Kozono et al. (52) (Figure 1A). This sequence is paired with the MHCIIβ native signal peptide to preserve targeting of the modified protein. Production of this transcript for use involves 1) preparing a plasmid encoding the transcript, 2) producing the transcript using *in vitro* transcription 3) encapsulating the transcript for immunization. All steps of this process can be accomplished with off-the-shelf commercial reagents, for optimal accessibility.

**Figure 1.**
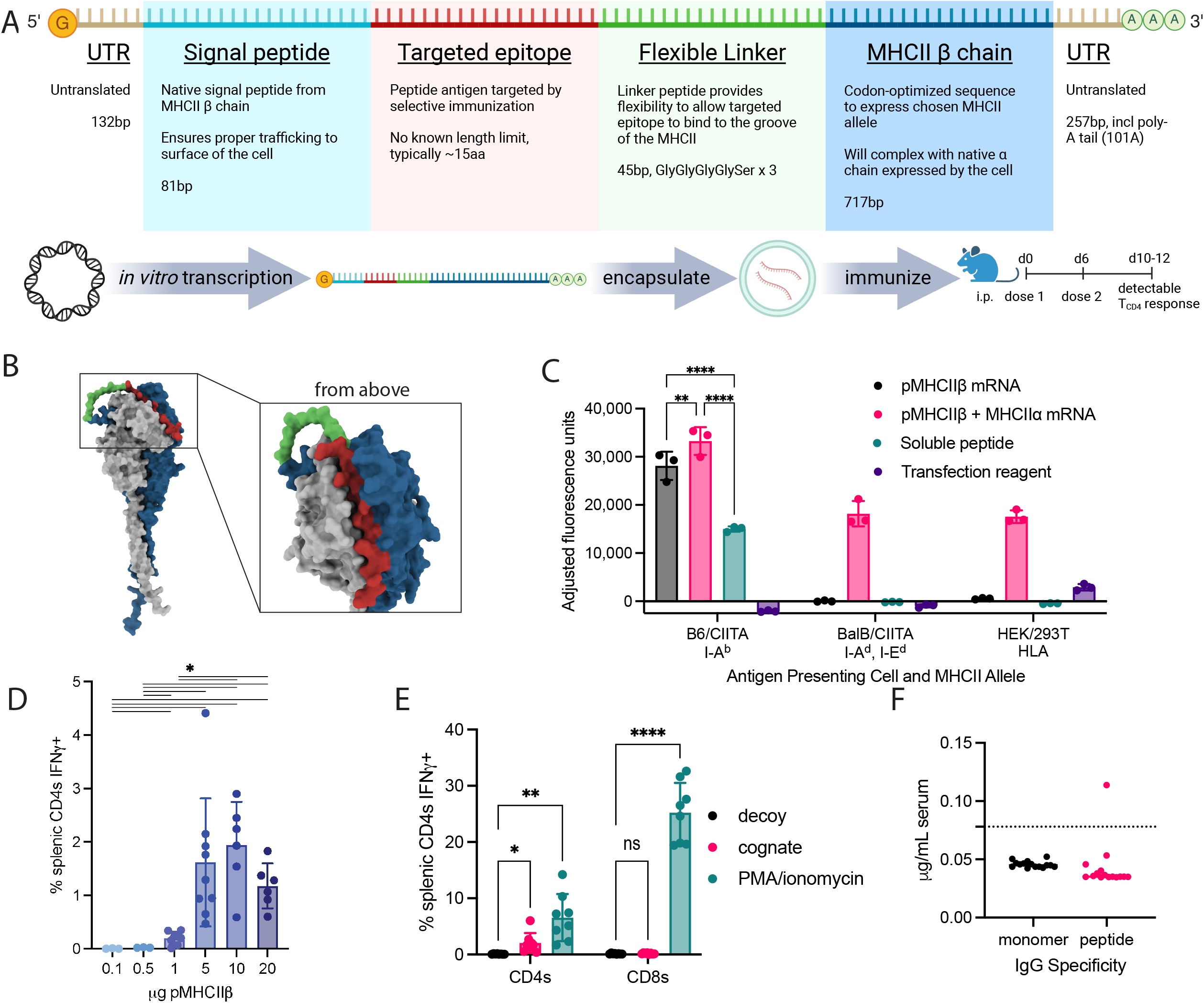
Custom mRNA platform encoding MHCIIβ with attached epitope allows efficient antigen presentation. (**A**) Schematic of mRNA transcript and mRNA production/use pipeline. (**B**) Colabfold model of pMHCIIb protein targeting Ova_323_/I-A^b^ complexing with WT MHCII a chain; targeted epitope in red, flexible linker in green, MHCIIb chain in blue, a chain in grey. (**C**) T hybridoma activation assay comparing activation of OTII reporter by cells transfected with Ova_323_/I-A^b^ pMHCIIβ +/-MHCII α, pulsed with Ova_323_ peptide, or treated with transfection reagent. Fluorescence values are normalized to media control, and analyzed via two-way ordinary ANOVA with Tukey’s multiple comparisons test with a single pooled variance. Representative of 3 independent experiments. (**D**) % of splenic CD4 T cells producing IFNγ in response to cognate peptide taken at d10 from mice immunized with a range of doses of pMHCIIβ. Analyzed via Brown-Forsythe and Welch ANOVA tests with Dunnett’s T3 multiple comparisons test. Data are drawn from 2 independent experiments, n=6. (**E**) % of splenic CD4^+^ T cells producing IFNγ in response to cognate peptide taken at d10 from mice immunized with 5μγ of pMHCIIβ mRNA, n=8. Analyzed via two-way ANOVA with Geisser-Greenhouse correction and Sidak’s multiple comparisons test. Data are drawn from 2 independent experiments. (**F**) ELISA of serum from d10 mice immunized with 5μγ of pMHCIIβ Ova323/IAb mRNA capturing IgG targeting Ova_323_/I-A^b^ monomer or Ova_323_ peptide. Data are drawn from 3 independent experiments, n=15. (**D-F**) each dot represents an individual mouse.

We hypothesized that this modified MHCII β chain would complex with native α chain to allow efficient presentation of the attached epitope. To evaluate this hypothesis, we first modeled the folding of these sequences using ColabFold v1.5.5 (Figure 1B) (53). We used a version of the transcript targeting the common model epitope from chicken ovalbumin, amino acids 323-339 (Ova_323_) presented on MHCII I-A^b^. The model predicted complexing of the two MHCII chains to form a normal binding groove, which was occupied by the Ova_323_ epitope. To test whether these proteins complexed as predicted and resulted in effective antigen presentation, we used a T_CD4_ hybridoma activation assay (Figure 1C). Briefly, cell lines were transfected with the transcripts encoding modified pMHCIIβ alone or in combination with WT α chain; cell lines were derived from C57Bl/6 and BALB/c backgrounds expressing WT MHCII, with HEK293T cells included as an additional negative control. Transfected cells were cocultured with an OTII T_CD4_ hybridoma reporter line for 18h. Addition of a fluorogenic reagent provides a readout of cognate antigen presentation. Results of this assay confirm that pMHCIIβ complexes with α chain to allow presentation of the attached epitope, including α chain produced endogenously by the cell. If a second transcript encoding the α chain is co-transfected into the cell, antigen presentation is possible even from a human cell line such as HEK293T, suggesting that expression of factors beyond the pMHCIIβ and α chain are not required. Notably, the antigen presentation from transfected cells appears to be very efficient, comparable to the addition of soluble peptide.

We next sought to determine the ideal dose of pMHCIIβ mRNA to use *in vivo* to provoke monospecific T_CD4_ responses. To do this, we immunized naïve C57Bl/6 mice intraperitoneally (i.p.) at d0 and d6 with a range of doses of pMHCIIβ mRNA (Figure 1D). At day 10, mice were euthanized and splenocytes were isolated and stimulated with soluble antigen overnight. The next day, cognate IFNγ responses were quantified via intracellular cytokine staining. This dose curve revealed a roughly logistic curve such that peak responses were achieved at 5ug and 10ug per dose. 5ug was selected as an ideal dose going forward; unless otherwise noted, this immunization schedule was used throughout. Notably, responses at 20ug appear to be trending slightly downward; this raises the question of not only diminishing returns, but possible inhibitory effects of higher doses.

In choosing this tool, we sought to create a platform for producing T_CD4_ responses in isolation, without provoking a T_CD8_ or antibody response. To measure T_CD8_ responses, we examined IFNγ responses in splenocytes from immunized mice (Figure 1E). We saw no significant increase in IFNγ production in the T_CD8_ compartment in response to cognate antigen, despite robust IFNγ signal in response to PMA/ionomycin, supporting our hypothesis that T cell responses to pMHCIIβ mRNA are confined to T_CD4_. To evaluate antibody responses, we collected postmortem serum from Ova_323_/I-A^b^ mRNA immunized mice and used ELISA to probe for IgG responses to either Ova_323_/I-A^b^ monomer or Ova_323_ soluble peptide (Figure 1F). At d10 post immunization, we found no detectable response to the monomer and only a single animal producing detectable response to soluble peptide. These results support the conclusion that the design of the pMHCIIβ immunogen sharply limits antibody responses.

### Immunization of mice with custom pMHCIIβ **mRNA elicits a consistent, polyfunctional Th1 response**

We next sought to determine the phenotype and properties of epitope-specific T_CD4_. Naïve mice were immunized i.p. with 5ug pMHCIIβ mRNA; splenocytes from immunized mice were cultured for 24h with either cognate or decoy soluble peptide. After incubation, culture media was collected, and secreted cytokines were identified and quantified via membrane-based antibody array (Figure 2A, S1). In examining cognate-specific cytokine responses, we noted a group associated with T_CD8_ recruitment (CCL3, CCL4)(54, 55)and with lymphocyte recruitment and T cell proliferation (CCL5, CXCL9, CXCL10).(56, 57) We also evaluated specific secretion of canonical Th1 cytokines (IL-2, TNFα, IFNγ) because T_CD4_ responses to mRNA vaccines often skew strongly toward this phenotype.(58, 59) Of these cytokines, only IL-2 was significantly higher with cognate antigen. However, we also noted significantly higher levels of the inflammatory cytokine IL-6. IL-6 is conventionally thought to skew T_CD4_ away from Th1, which would be inconsistent with the previously noted IL-2 production.(60, 61) However, IL-6 also supports Tfh differentiation, a process efficiently induced by mRNA vaccination.(5, 7, 62, 63) Finally, we observed a significant increase in soluble CD54 (ICAM-1), a signal associated with a wide variety of inflammatory states.(64–67) In addition to these cognate-specific signals, we observed substantial secretion of IL-16 in response to both cognate and decoy antigen, most likely in response to anti-CD28 antibody included in the coculture.(68)

**Figure 2.**
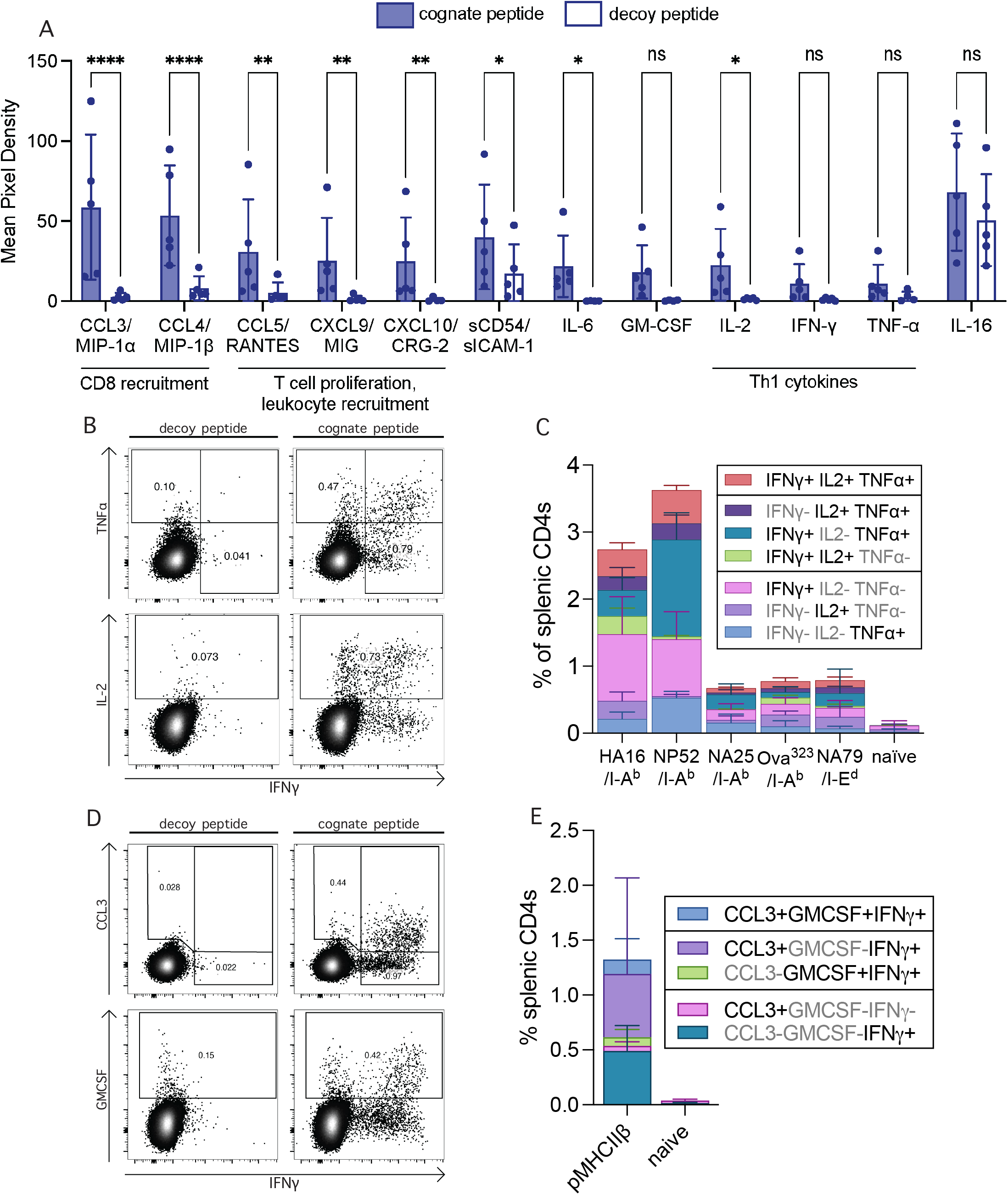
Immunization with pMHCIIb mRNA provokes polyfunctional Th1 responses. (**A**) Select results of cytokine detection array. d10 splenocytes were taken from mice immunized with pMHCIIβ Ova323/IAb mRNA, cocultured with Ova323 or decoy peptide, cytokines were measured from culture media. Each dot represents a mouse, n=5. Analyzed via mixed-effects model, with Sidak’s multiple comparisons test. (**B,D**) Representative flow cytometry plots highlighting cyto-kine production in response to cognate peptide by CD4 T cells from pMHCII β immunized mice. (**C,E**) Quantification of cytokine production in response to cognate peptide by CD4 T cells from pMHCII β immunized mice, n=6-9 per group, drawn from 2-3 independent experiments.

We used intracellular cytokine staining to further explore the functionality of the T_CD4_ population elicited by the selective immunization. We first evaluated responses of the Th1 panel of IFNγ, TNFα, and IL2 in response to a panel of pMHCIIβ mRNA targeted at different influenza epitopes and MHCII alleles (Figure 2B-C, Table 1, S2A). The results of this analysis revealed that pMHCIIβ mRNA consistently produces a polyfunctional Th1 response, although the magnitude of the response varies, potentially due to differences in precursor frequency. We next examined expression of two additional cytokines, CCL3 and GM-CSF, finding both were produced by IFNγ+ cells in response to cognate antigen (Figure 2D-E, Figure S2B). Our results demonstrate that our platform produces a robust, polyfunctional Th1 response to a variety of epitope/MHCII pairings.

| Epitope | Protein of origin | Species | MHCII allele | AA primary sequence |
| --- | --- | --- | --- | --- |
| Ova323 | Ovalbumin | Chicken | I-Ab | ISQAVHAAHAEINEAGR |
| HA16 | Hemagglutinin | Influenza A |  | RSWSYIVETPNSENGIC |
| NA25 | Neuraminidase |  |  | GDV FVIREPFISCSH |
| NP52 | Nucleoprotein |  |  | LLQTSQVYSLIRPNENP |
| NA79 | Neuraminidase |  | I-Ed | IRGWAIYSKDNSIRI |

### Immunization of mice with custom pMHCIIβ **mRNA prevents bacterial shedding in a model of Salmonella enterica infection**

We next wanted to evaluate the *in vivo* impact of T_CD4_ elicited by pMHCIIβ mRNA. We selected *Salmonella enterica spp typhymurium* (STm) as our infection model; immune responses to STm are highly dependent upon T_CD4_ responses, especially Th1 responses, as demonstrated in depletion and knockout experiments.(23, 69, 70) To test the effect of T_CD4_ provoked by our pMHCIIβ mRNA on the immune response to STm, we used STm expressing Ova_323_ antigen to infect C57Bl/6 mice immunized with pMHCIIβ mRNA targeting Ova_323_ or a decoy antigen (Figure 3). Stool samples were collected from mice to evaluate levels of bacterial colonization throughout the infection. At d4 post infection, we observed a significantly lower level of colonization of the mice immunized with the on-target pMHCIIβ, supporting our hypothesis that T_CD4_ produced by our platform function within the immunized animal as part of a larger immune response.

**Figure 3.**
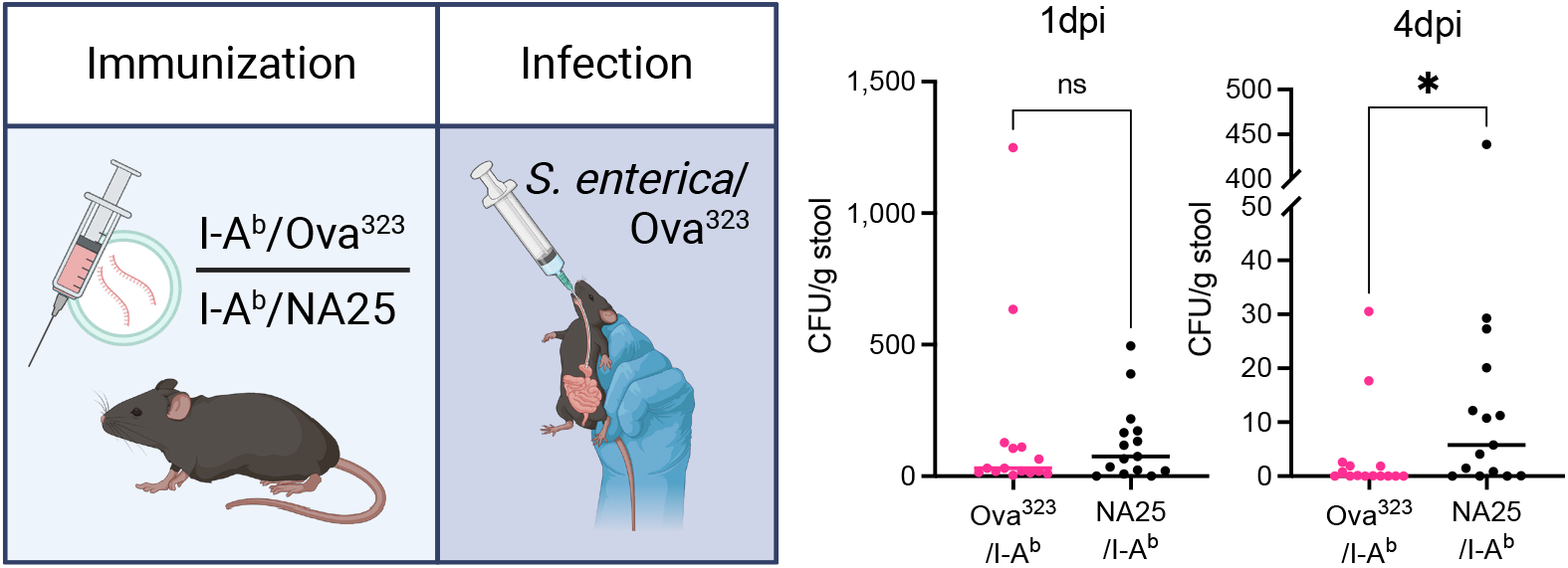
CD4 T cells elicited by pMHCIIb mRNA limits Salmonella infection. Mice were immunized with pMHCIIβ mRNA targeting Ova323/I-Ab or NA25/I-Ab, then infected with *S. enterica spp. typhimurium* strain SL7207 expressing Ova323. Stool from each mouse were collected and cultured to determine bacterial burden. Each dot represents an individual mouse; data are drawn from 3 independent experiments, n=15 per group. Analyzed via Mann-Whitney test, \**p*<0.05

In conclusion, we have developed a novel method using a customizable mRNA platform to produce robust, isolated, and monospecific T_CD4_ responses exhibiting a strongly polyfunctional Th1 phenotype and *in vivo* activity.

## Acknowledgements

We thank the Children’s Hospital of Philadelphia flow cytometry core and animal husbandry staff. We thank Dieter Schifferli of the University of Pennsylvania School of Veterinary Medicine for his kind gift of recombinant Salmonella. M.J.H. was supported in part by the Cancer Research Institute as a Cancer Research Institute

Irvington Fellow and by the Roberts Family–Katalin Karikó Fellowship in Vaccine Development from the Aileen K. and Brian L. Roberts Family Foundation via the University of Pennsylvania Institute for Immunology & Immune Health (I3H).

## Disclosures

The authors have no financial conflicts of interest.

## Notes

### Competing Interest Statement

The authors have declared no competing interest.

