## Supplemental Figure 1 for "Cutting Edge: An mRNA Platform to Create Isolated, Monospecific Th1 Responses"

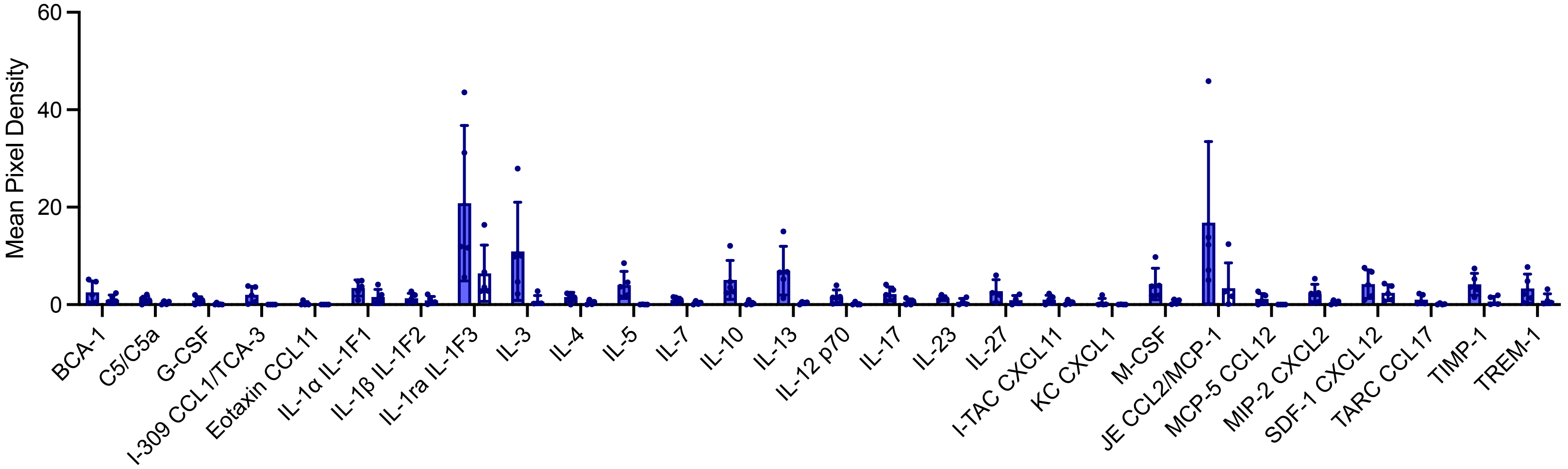


**Figure S1: Additional results of cytokine array in Figure 2A** d10 splenocytes were taken from mice immunized with pMHCIIβ $Ova_{323}$/I-$A^b$ mRNA, cocultured with $Ova_{323}$ or decoy peptide, cytokines were measured from culture media. Each dot represents a mouse, n=5. Analyzed via mixed-effects model, with Sidak's multiple comparisons test.
