## Supplemental Figure 2 for "Cutting Edge: An mRNA Platform to Create Isolated, Monospecific Th1 Responses"

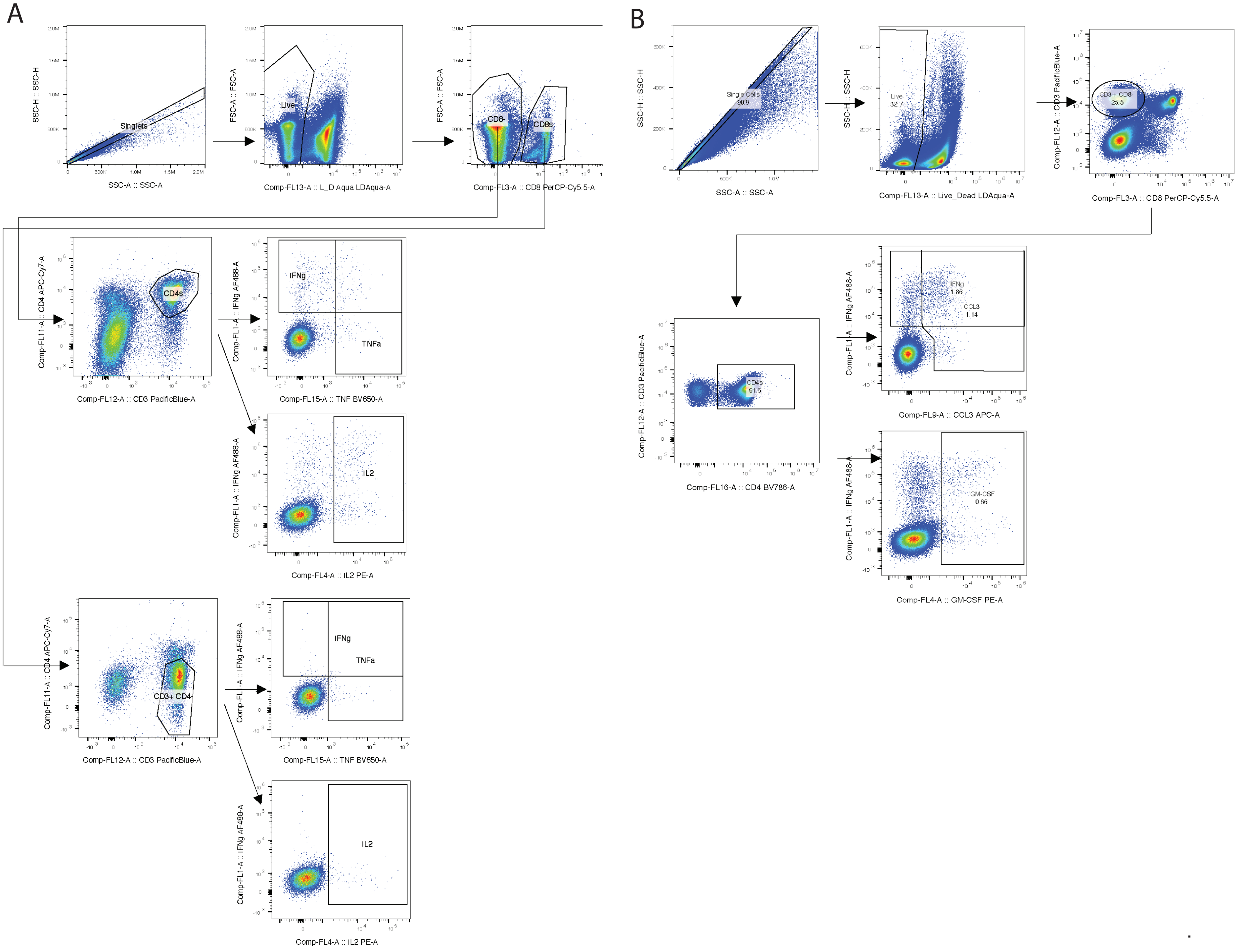


Figure S2 | Figure 2 gating strategies (A) Gating strategies for Figure 2B,C (B) Gating strategies for Figure 2D,E.
